# Evaluating user experience with immersive technology in simulation-based education: a modified Delphi study with qualitative analysis

**DOI:** 10.1101/2022.09.26.509545

**Authors:** Chris Jacobs, Georgia Foote, Michael Williams

## Abstract

**Background:** Immersive technology is becoming more widespread in simulation-based medical education with applications that both supplement and replace traditional teaching methods. There is a lack of validated measures that capture user experience to inform of the technology utility. We aimed to establish a consensus of items and domains that different simulation experts would include in a measure for immersive technology use.

**Methods:** A 3-stage modified Delphi using online software was conducted to support the conceptual framework for the proposed measure. The first round was informed by prior work on immersive technology in simulation. In the first round, participants were asked to describe what we could measure in simulation-based education and technology. Thematic analysis generated key themes that were presented to the participants in the second round. Ranking of importance in round 2 was determined by mean rank scores. The final round was an online meeting for final consensus discussion and most important domains by experts were considered.

**Results:** A total of 16 simulation experts participated in the study. A consensus was reached on the ideal measure in immersive technology simulation that would be a user questionnaire and domains of interest would be: what was learnt, the degree of immersion experienced, fidelity provided, debrief, psychological safety and patient safety. No consensus was reached with the barriers that this technology introduces in education.

**Conclusions:** There is varied opinion on what we should prioritise in measuring the experience in simulation practice. Importantly, this study identified key areas that aids our understanding on how we can measure new technology in educational settings. Synthesising these results in to a multidomain instrument requires a systematic approach to testing in future research

## Background

Teaching through simulation is well established in medical disciplines.(1) The pedagogy underlying simulation is based on deliberate practice through recreation of clinical scenarios, irrespective of the enabling technology. An educational experience is constructed with elements that resemble relevant environments, with a greater or lesser degree of fidelity, and match the functional task with the learner’s engagement.(2) For example, we might describe a high-fidelity manikin and clinically accurate context, for training for students in basic life support.(3)

In recent decades there are examples of technological innovation enhancing educational effectiveness through increased immersion, associated with improved motivational states.(4) There are many technological modalities of simulation, summarised in table 1.

**Table 1.**
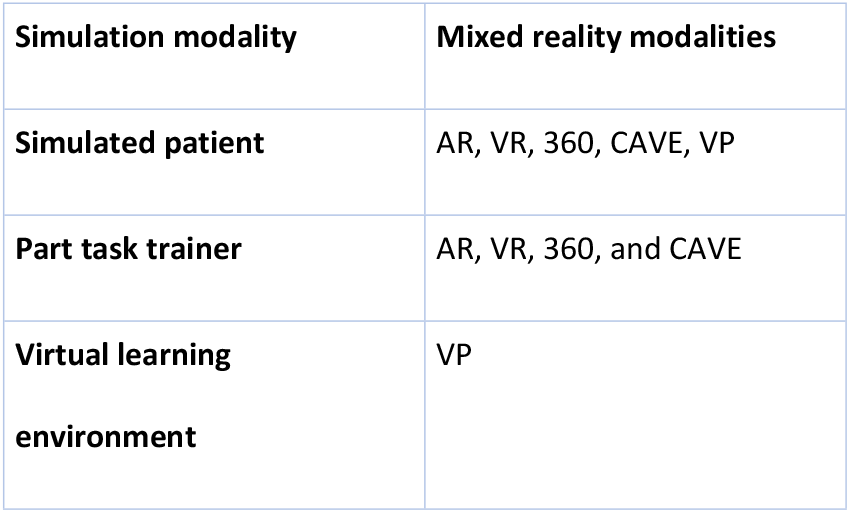
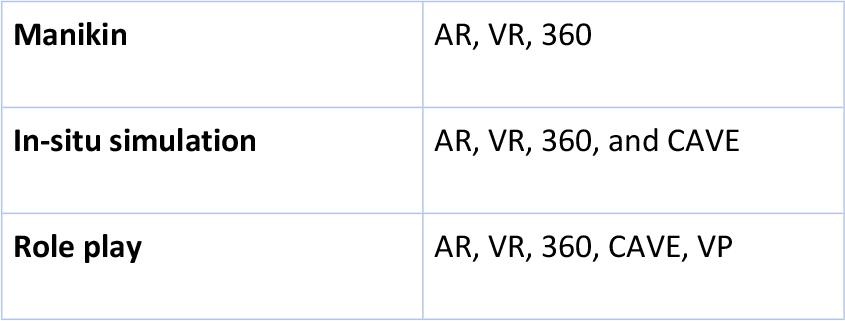
Simulation modalities with mixed reality methods. AR- Augmented reality, VR- Virtual Reality, 360-360-degree video, CAVE- Cave Automatic Virtual Environments, VP- Virtual patient. Modified from Forrest and McKimm (5)

| Simulation modality | Mixed reality modalities |
| --- | --- |
| Simulated patient | AR, VR, 360, CAVE, VP |
| Part task trainer | AR, VR, 360, and CAVE |
| Virtual learning environment | VP |
| **Manikin** | AR, VR, 360 |
| **In-situ simulation** | AR, VR, 360, and CAVE |
| **Role play** | AR, VR, 360, CAVE, VP |

The Simulation Process provides an overarching structure to simulation-based education (SBE) (figure 1). Immersive technology integration in SBE requires consideration of all levels of the process, as no technological modality is inherently superior, and each possesses different strengths and weaknesses relevant for different scenarios.

**Figure 1.**
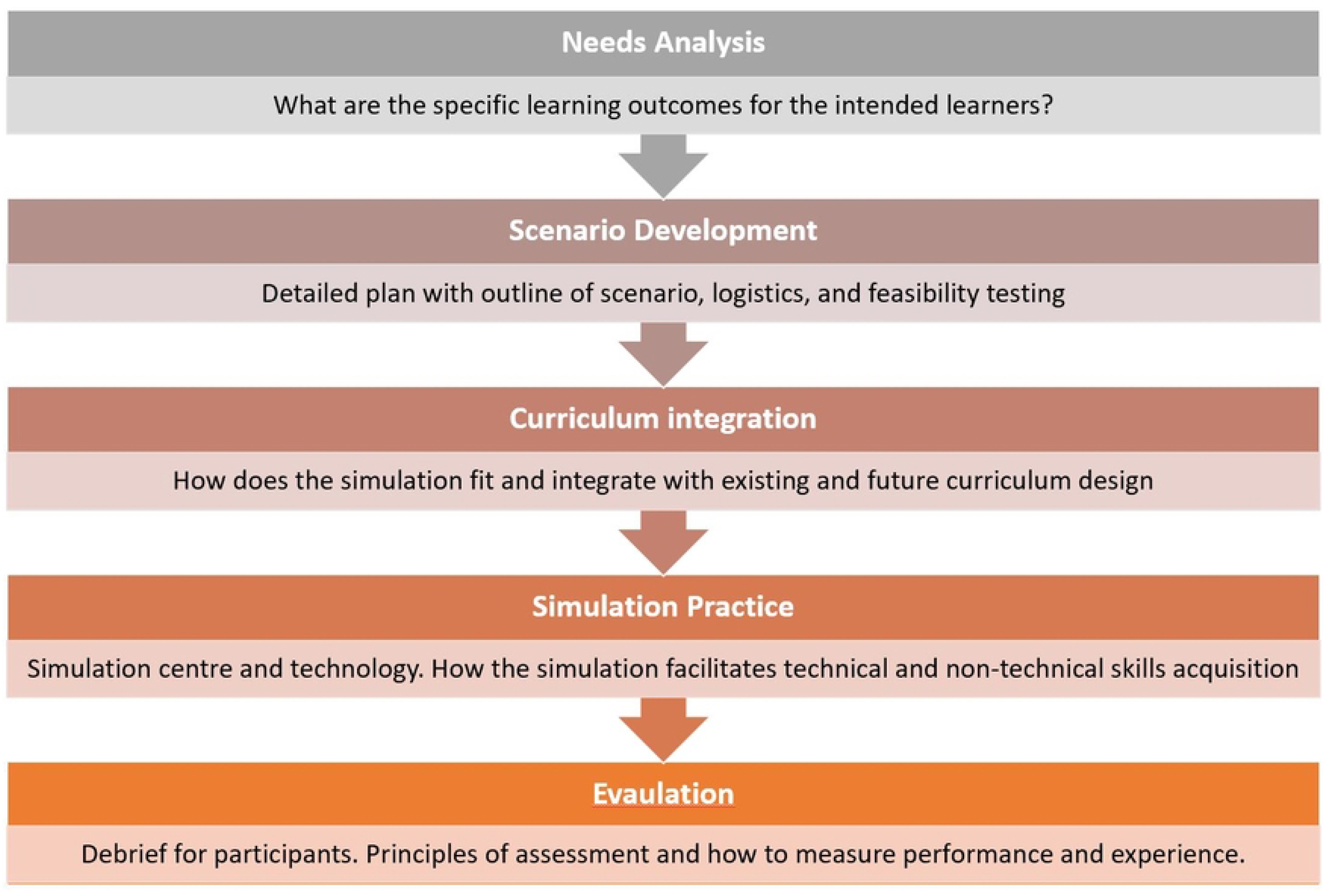
Simulation process design in simulation-based education

The reality-virtuality continuum describes a range of simulation environments, from real world to virtual world. Within the continuum, the sensory stimuli can be a mix of real and virtual objects. For example, Mixed Reality (MR) is an overarching heading that includes augmentation of real-life (AR), and virtual environments in a head-mounted display (VR).(6) The extent of real-world disconnect and suspension of disbelief is dependent on the technology. Coherence reflects the authenticity or fidelity with which the technology matches to the real-world, and again there is a spectrum of coherence.(7)

There is ever growing evidence that immersive technology benefits healthcare students and qualified health professionals in simulation practice.(4, 8–10) Studies investigating the effectiveness of differing technologies in varied medical specialties is increasing year on year, with research in both VR and AR doubling in the period of 2020-2021.(11)

In a systematic review on immersive technology in healthcare education, over 50 methods were described of measuring healthcare practice, which could be broadly separated into: cognitive objective, procedural skills objective, or subjective behavioural. Capturing this information helps educators judge the performance of participants and the utility of the technology. The quality of the assessment tool used is essential for evaluating how potentially generalisable the results are (12, 13). However, shortcomings in the validity and evidence supporting the use of many assessment tools in SBE exist.(13, 14)

The Delphi technique is widely accepted as an effective group-based exercise, which aims to integrate multiple interpretations and opinions on a given topic.(15, 16) It has 4 defining characteristics: group anonymity, repeating questions (iteration), feedback to participants, and statistical description of responses.(17) The Delphi researchers orchestrate focal issues surrounding the mode of delivery, the threshold for consensus and encouragement of responses.(15) Delphi techniques supplement analyses of published literature on a topic in preparing a comprehensive framework.

The overarching aim of this project is to create a *de* novo brief, valid and reliable multi-dimensional measure that will inform educators and learners on the users’ experiences of technology in simulation practice. The domains that could be considered important in quantifying a user’s experience are yet not fully understood. The Delphi approach allows for a diverse number of experts in the field of SBE to be brought together to address this.

## Materials and Methods

The study was funded by Health Education England (HEE) Southwest Simulation Network as part of a 2-year project collaboration with University of Bath and simulation network. This study will establish a consensus for generating a new measure’s theoretical and conceptual framework, that can be further examined with cognitive interviewing and quantitative exploration in future work.

### Modified Delphi method

A steering group consisting of authors (CJ, GF, and MW) guided three-stages of convergent opinion and consensus building(18) between May and August 2022. The conventional Delphi method uses iterative rounds of data collection from groups of experts, with subsequent feedback on responses, with the opportunity for them to rethink their response until consensus is met.(19) Online questionnaire collection facilitates this process for response collection and feedback, in short time frames.(20)

A hybrid of the conventional method, a modified Delphi method was adopted, which collects initial participant opinions online, followed by in-person group meetings for final consensus discussion.(15, 21) An *a priori* response rate was set at 70% for Delphi round 2, which was deemed as minimally acceptable to minimise bias.(22) A lower response to the online meeting was deemed acceptable as it required greater commitment, and drop-out linked to additional demands of attending.(23) Figure 2 flow chart illustrates the Delphi process, and a reporting checklist is provided in supplementary material.

**Figure 2.**
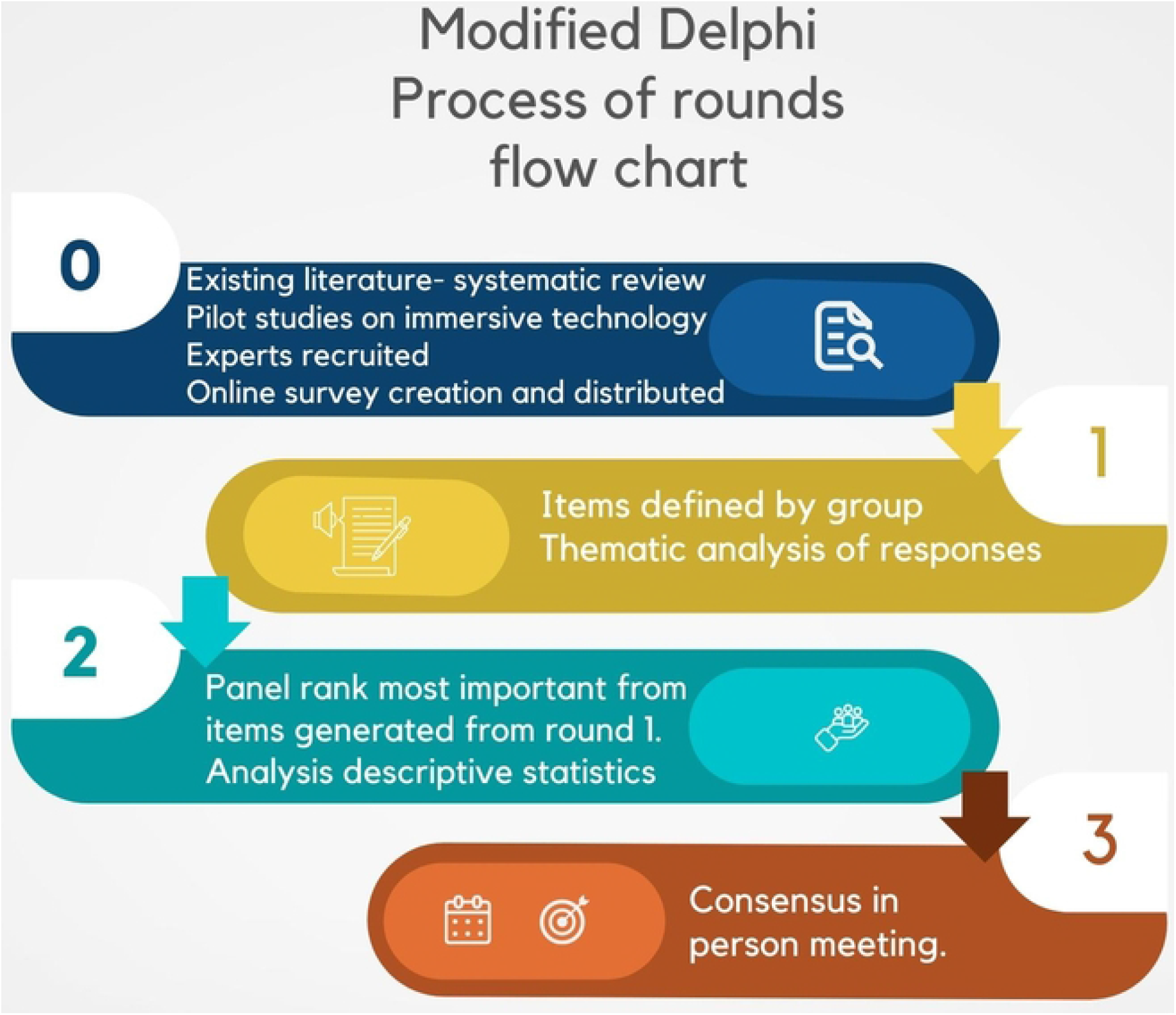
Evaluating user experience with immersive technology in simulation-based education Modified Delphi process flow chart

### Participants

The target population for this study will be individuals engaged in SBE. Stake holders included doctors, technicians, managers, and administrators involved in simulation for healthcare education (table 2). Purposive sampling from a targeted group allowed creation of a group of experts, collectively offering a range of experience in healthcare simulation. It was the aim to recruit 10 to 20, which has been described as the panel size to allow meaningful statistical analysis.(20, 24) Recruitment was via a Southwest simulation network email list as a single advert.

**Table 2.** Eligibility criteria- key inclusion and exclusion criteria.

|  |
| --- |
| Key inclusion criteria |
| Experience with simulation in healthcare education setting in the Southwest of England |
| Able to access the internet (Study sign-up and intervention could only be accessed through the internet). |
| Willing to participate in 2 rounds of questionnaires AND/OR willing participate in a virtual meeting. |

### Data collection and analysis

#### Delphi Round 1

This was an online questionnaire. Questionnaire design and distribution was done using Jisc Online surveys (https://www.onlinesurveys.ac.uk accessed September 2022), software which meets data protection regulations. Participants’ information and consent were embedded in the survey prior to questions. The Delphi study was approved by Health Research Authority (research ethics committee reference 22/PR/0339 and integrated research application system project ID 312830). Research sponsor was the Great Western Hospital NHS Foundation Trust (UK).

Participants’ age and roles were collected to inform authors of range of their simulation experience. Otherwise, question development was drawn from 2 sources: firstly, a qualitative study exploring themes of participant experience in Virtual Reality within an education setting,(25) and secondly, a systematic review was undertaken on immersive technology in healthcare education covering literature from 2002 and 2022.(11) In the systematic review, data from 246 papers on learning theory and on measures adopted in simulation practice with immersive technology were analysed. Outcomes from both these sources were discussed between authors CJ and MW to inform questionnaire design and participant information sheets. The questions focused on the modalities of learning in SBE, how this might be measured, and what potential barriers to use of emergent technology.

Round 1 provided qualitative data: panelists responded with free text answers in the form of a non-prioritised list. Thematic analysis was conducted to these open-ended questions with a 6-step process.(26, 27) Inductive reasoning guided the analysis of responses to identify key patterns and relationships. Generation of initial codes was done independently by CJ and GF. Further interpretative analysis was undertaken by CJ and GF, specifically repeated conversations and code sorting to identify overarching themes, that were eventually defined in short phrases or words that unified the narrative of responses. A thematic map was created as a pictorial summary for feedback to the expert panelists.

#### Delphi Round 2

This was an online questionnaire. Those participating in round 1 were invited to respond to round 2, which included the same questions with the newly identified and defined key themes resulting from round 1 as options for ranking items in importance from highest to lowest using a check-list response. Means, median and interquartile range (IQR) were calculated using StatsDirect version 3.3.5 for the ordinal round 2 responses.(24) Statistical significance of data was determined as p <0.05, assessed using non-parametric Friedman’s test and weighted Kappa to describe strength of agreement between experts ranking.(28) Kappa values were treated as 0.00 to 0.10 conferring poor agreement, 0.11 to 0.20 conferring slight agreement, 0.21 to 0.40 conferring fair agreement, 0.41 to 0.60 conferring moderate agreement, and 0.61 to 0.80 conferring substantial agreement.(29) Mean rankings were complemented with a boxplot visual summary for each question(15), and overarching thematic maps from Round 1 were distributed to participants prior to round 3.

#### Delphi Round 3

The online meeting was held using Zoom. The meeting was chaired by CJ and field notes collated by GF. Revealing of majority opinions prior to the final meeting enabled experts to reflect on group thinking with potential disagreement amongst experts during Round 2 seen, although, consensus was the ultimate aim. In this round sharing of opinions was promoted, and a discussion facilitated clarification of any misunderstandings (30). Chairing of a meeting promotes encouragement of those attending and avoiding dominance of voices.(31) An effort was made to have everyone contribute to the conversations. Consensus criteria, set before round 3, for identification of the top 4 themes from responses to questions 1 and 2 was set at 75%, and for question 4, the top single method of measuring was set at 75% of those attending the meeting(32) with opinions collected using the Zoom polling tool.

## Results

There were 16 participants recruited from an email sent to 350 on a mailing list. True response rate in relation to inclusion and exclusion criteria could not be calculated. Table 3 shows the demography of the group in this study.

**Table 3.** Age and roles of participants. SBE- Simulation based education.

| SBE role | Age |
| --- | --- |
| Outreach simulation co-ordinator | 33 |
| Clinical Teaching Fellow | 26 |
| Simulation Medical Lead | 37 |
| Programme Lead for Simulation & TEL for HEE southeast | 48 |
| Paediatric Simulation fellow at University Hospitals Bristol and Weston NHS Foundation Trust | 33 |
| Nurse Education (university) | 46 |
| Simulation Facilitator | 33 |
| Primary Care Simulation Fellow | 38 |
| Clinical Skills Tutor to final Year Medical Students | 53 |
| Resuscitation & Simulation Officer | 28 |
| GP tutor | 33 |
| Previous clinical teaching fellow | 32 |
| Clinical teaching fellow | 30 |
| Senior Technical Instructor | 44 |
| Simulation and Clinical Skills Technician | 46 |

### Qualitative Round 1

Questions asked and key themes emerging from round 1 are seen presented in Figure 3 below.

**Figure 3.**
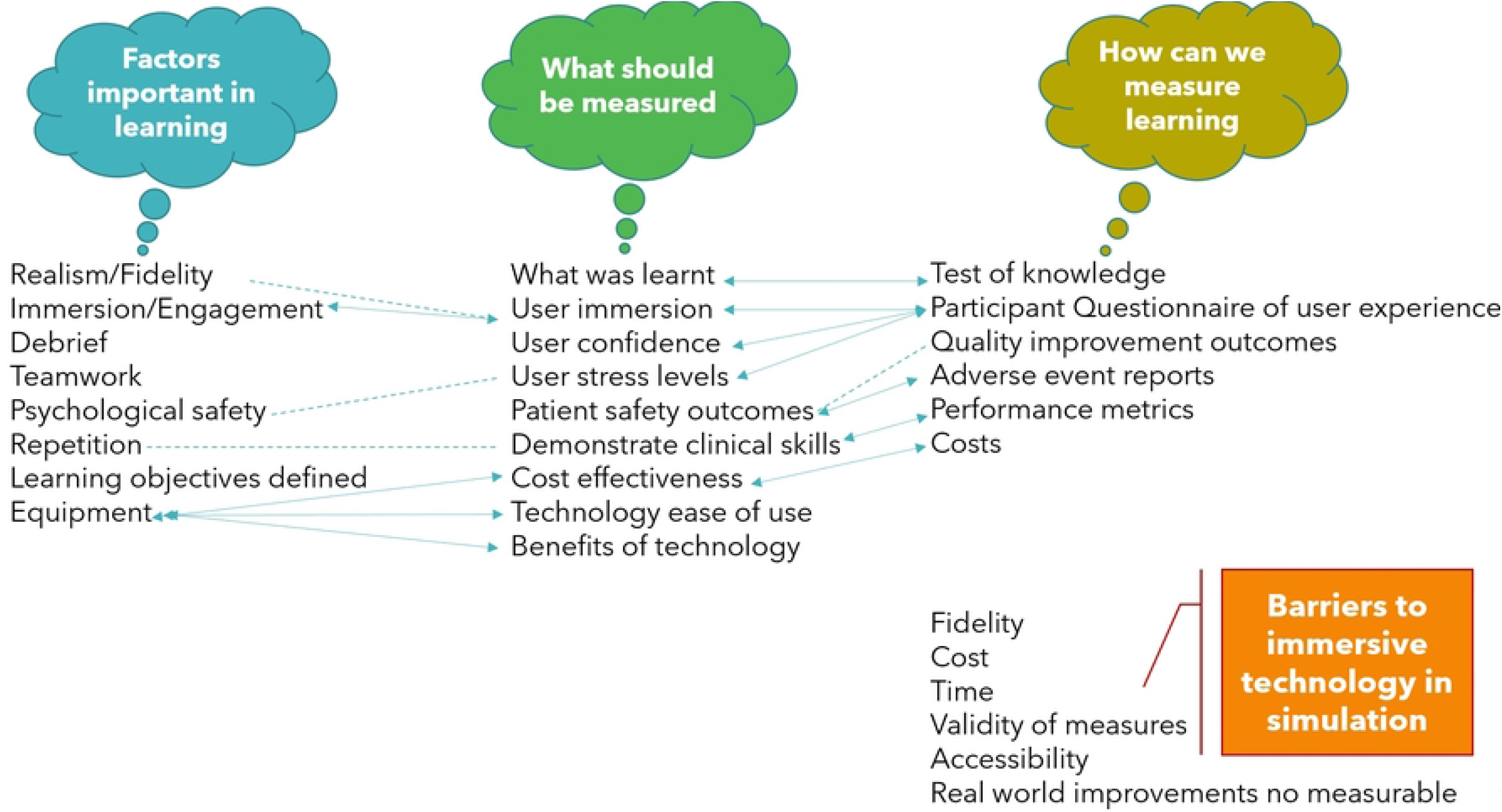
Thematic map with the 4 questions abbreviated. Arrows indicate a 2-way link of response between questions. Dotted line indicates association of responses. Red line represents barriers.

### Quantitative Round 2

Thirteen panelists from initial 16 recruited (81%) responded to round 2.

The 4 questions were individually analysed. Figure 4 panel A-D are boxplots with medians and IQR for each question

**Figure 4.**
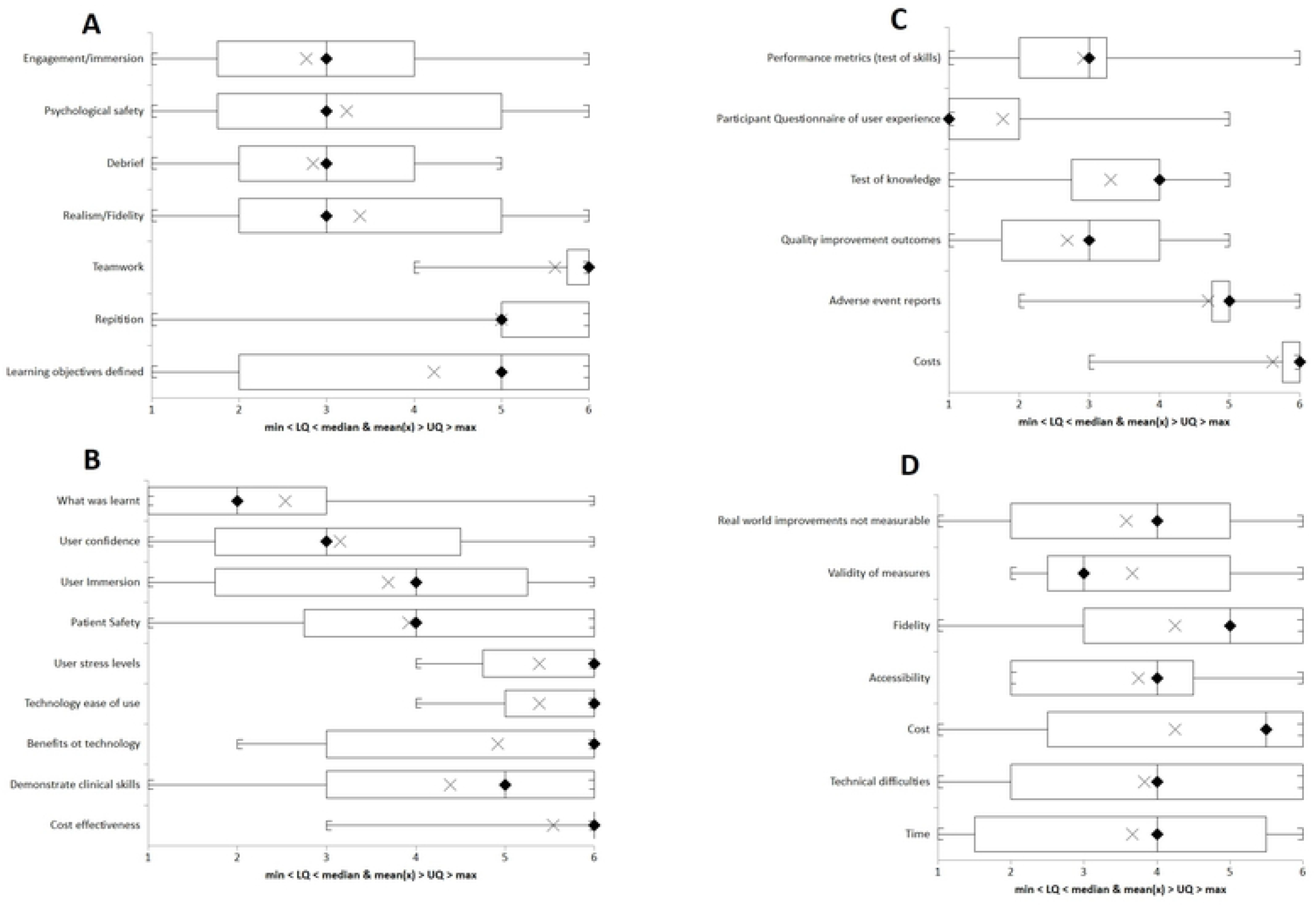
Boxplot of theme rankings. A- Question 1, B-Question 2, C-Question 3, and D- Question 4. Key, LQ-lower quartile, UQ- Upper quartile, cross marks mean, rhombus marks median.

### Question 1- List up to 5 factors that you think are important in learning in simulation?

Top four ranked responses in order: Engagement/immersion (mean rank 2.76), Debrief (mean rank 2.85), psychological safety (mean rank 3.23), and realism/fidelity (mean rank 3.39). Table 4 summarises the descriptive data for question 1.

**Table 4.** Descriptive results for question 1 with mean ranking.

| Themes | Engagement<br>Immersion | Psychological<br>safety | Debrief | Realism/Fid<br>elity | Teamwork | Repetition | Learning<br>objectives<br>defined |
| --- | --- | --- | --- | --- | --- | --- | --- |
| Valid data | 13 | 13 | 13 | 13 | 13 | 13 | 13 |
| Missing<br>data | 0 | 0 | 0 | 0 | 0 | 0 | 0 |
| Mean | 2.76 | 3.23 | 2.85 | 3.39 | 5.62 | 5.00 | 4.15 |
| Upper<br>quartile | 4 | 5 | 4 | 5 | 6 | 6 | 6 |
| Median | 3 | 3 | 3 | 3 | 6 | 5 | 5 |
| Lower<br>quartile | 2 | 2 | 2 | 2 | 6 | 5 | 2 |
| IQR | 2 | 3 | 2 | 3 | 0 | 1 | 4 |

Friedman’s test indicated rank patterns (*p* <0.001) with a degree of difference in ranking.

Weighted Kappa 0.21 (*p*<0.001) indicating fair agreement in rankings between participants.

### Question 2- List what you think should be measured

Top four ranked responses in order: What was learnt (mean rank 2.54), user confidence (mean rank 3.15), user immersion (mean rank 3.69), and patient safety (mean rank 3.92). Table 5 summarises the descriptive data for question 2.

**Table 5.** Descriptive results for question 2 with mean ranking

| Themes | What was learnt | User confidence | User Immersion | Cost effectiveness | Demonstrate clinical skills | Benefits of technology | Technology ease of use | User stress levels | Patient Safety |
| --- | --- | --- | --- | --- | --- | --- | --- | --- | --- |
| Valid data | 13 | 13 | 13 | 13 | 13 | 13 | 13 | 13 | 13 |
| Missing data | 0 | 0 | 0 | 0 | 0 | 0 | 0 | 0 | 0 |
| Mean | 2.54 | 3.15 | 3.69 | 5.54 | 4.38 | 4.92 | 5.38 | 5.38 | 3.92 |
| Upper quartile | 2 | 4 | 5 | 6 | 6 | 6 | 6 | 6 | 6 |
| Median | 2 | 3 | 4 | 6 | 5 | 6 | 6 | 6 | 4 |
| Lower<br>quartile | 1 | 2 | 2 | 6 | 3 | 3 | 5 | 5 | 3 |
| Interqua<br>rtile<br>range | 1 | 2 | 3 | 0 | 3 | 3 | 1 | 1 | 3 |

Friedman’s test indicated rank patterns (*p* <0.001) with a degree of difference in ranking.

Weighted Kappa 0.22 (*p*=0.0003) indicating a fair agreement in rankings between participants.

### Question 3- What measurement methods could you use?

Top four ranked responses in order: Participant questionnaire of user experience (mean rank 1.77), quality improvement outcomes (mean rank 2.69), performance metrics (mean rank 2.92), and test of knowledge (mean rank 3.31). Table 6 summarises the descriptive data for question 3.

**Table 6.** Descriptive results for question 3 with mean ranking

| Themes | Performance<br>metrics (test<br>of skills) | Participant<br>questionnaire of<br>user experience | Test of<br>Knowledge | Quality<br>improvement<br>outcomes | Adverse event<br>reports | Costs |
| --- | --- | --- | --- | --- | --- | --- |
| Valid data | 13 | 13 | 13 | 13 | 13 | 13 |
| Missing data | 0 | 0 | 0 | 0 | 0 | 0 |
| Mean | 2.92 | 1.77 | 3.31 | 2.69 | 4.69 | 5.62 |
| Upper quartile | 3 | 2 | 4 | 4 | 5 | 6 |
| Median | 3 | 1 | 4 | 3 | 5 | 6 |
| Lower quartile | 2 | 1 | 3 | 2 | 5 | 6 |
| Interquartile range | 1 | 1 | 1 | 2 | 0 | 0 |

Friedman’s test indicated rank patterns (*p* <0.001) with a degree of difference in ranking.

Weighted Kappa 0.22 (*p* <0.001) indicating a fair agreement in rankings between participants.

### Question 4- List the barriers you consider in “measuring immersive technology” in simulation and medical education

Top four ranked responses in order: Real world improvements not measurable (mean rank 3.58), validity of measures (mean rank 3.67), cost (mean rank 3.67), and technical difficulties (mean rank 3.83). Table 7 summarises the descriptive data for question 4.

**Table 7.** Descriptive results for question 4 with mean ranking

| Themes | Real world improvements not measurable | Validity of measures | Fidelity | Accessibility | Cost | Technical difficulties | Cost |
| --- | --- | --- | --- | --- | --- | --- | --- |
| Valid data | 13 | 13 | 13 | 13 | 13 | 13 | 13 |
| Missing data | 0 | 0 | 0 | 0 | 0 | 0 | 0 |
| Mean | 3.58 | 3.67 | 4.25 | 3.75 | 4.25 | 3.83 | 3.67 |
| Upper quartile | 5 | 5 | 6 | 5 | 6 | 6 | 5 |
| Median | 4 | 3 | 5 | 4 | 5 | 4 | 4 |
| Lower quartile | 2 | 2 | 3 | 2 | 2 | 2 | 1 |
| Interquartile range | 3 | 2 | 3 | 2 | 3 | 4 | 4 |

Friedman’s test indicated no significant rank patterns (*p*=0.97).

#### Delphi Round 3

All participants of both rounds 1 and 2 were invited to join a virtual meeting hosted on Zoom. Five participants (36%) attended: 2 doctors, 2 nurses, and 1 simulation technician. There was a short presentation with a brief recap on literature surrounding immersive technology in medical education and each question was discussed in turn with results from round 2 available to prompt discussion. Table 8 shows the consensus reached for each of the 4 questions. The lack of spread to data in question 4 meant it was not appropriate for a consensus as each held equal weight.

**Table 8-.**

| Question | Consensus |
| --- | --- |
| List up to 5 factors that you think are important in learning in simulation? | 100% agreement with:<br>Engagement/Immersion<br>Psychological safety<br>Debrief<br>Realism/Fidelity |
| List what you think should be measured | 60% agreement with:<br>What was learnt<br>User confidence<br>User Immersion<br>Patient Safety<br>100% agreement if Cost effectiveness is added to above. |
| What measurement methods could you use? | 100% agreement with:<br>Participant questionnaire of user experience |
| List the barriers you consider in “measuring immersive technology” in simulation and medical education | No consensus reached |

Field notes taken during the final meeting created discussion points with each question presented. Although, no further thematic analysis was undertaken, selected quotations highlighted the expert opinion on the way we might learn from immersive technology enhanced learning (TEL) and how we might measure it. Complete field notes are available in supplementary material.

Participant J (doctor) in response to question 1.

> *“I think, fidelity is high up – with enjoyment and immersion, then debrief”*

Following this statement, participant DM (simulation technician) echoed J thoughts.

> *“engagement is instantaneous feedback about simulation…then can consolidate learning with debrief”*

Participant SK (doctor) reflecting on question 2.

> *“…simulation as a concept can be a challenge…cost effectiveness is hard to measure but is very important to collect data on”*

Following this participant J responded to SK in question 2.

> *“ This tech is novel and is useful compared to standard methods – so immersion and testing how useful the technology is important”*

Expert consensus was quickly reached with question 3 as acceptability on measuring user experience was the most important, although it was acknowledged that this might not capture all relevant factors.

> *“yes and no – good as it is easy to ask what they think – but whether the tech has made a difference after the session could be measured… difficult to measure maybe best measure is quality improvement and adverse events”*

## Discussion

Key aspects of relevant domains on measuring user experience of immersive technology in simulation were identified using the modified Delphi method in this study, with a consistency of responses evident across a heterogeneous groups of experts. It was agreed that a measure ideally would collect information on: what was learnt, the degree of immersion experienced, fidelity provided, debrief, psychological safety and patient safety. Additionally, a technological assessment should include a cost-effectiveness evaluation. There was 100% consensus in round 3 that a psychometric measure of user experience using a participant questionnaire would be the most suitable format to explore use of immersive technology in SBE.

There are numerous measures, subjective and objective, in existence for assessing a particular immersive technology in a certain situation.(33) These are often created for the purpose of the individual study in which they are used, without any argument for their validity.(13) Furthermore, there is a risk that instruments used to collect data become obsolete as emergent technology surpasses existent training methods. An international Delphi study was conducted to ensure researchers shared definitions of and terminology about instrument properties.(34) This study clarified what was meant by key measurement properties: reliability, validity, responsiveness and interpretability. These properties need to be considered when any outcome measurement is tested. The findings of a robust literature review supported by this Delphi study have led to a conceptual model of how the utility of immersive technology in healthcare education can be measured, indeed what to measure. This will inform development of a new evidence-based measure, which can then undergo field testing with various technologies and settings, that will enable the properties to be tested and the measure refined. The process of validity is not a single task, but testing hypotheses in a continuous process to see whether scores are consistent with the model intended.(35)

Although, various methods of Delphi are possible, a qualitative first round is useful as it allows experts to contribute ideas beyond current established or published knowledge. Reliability and validity of the Delphi study may be improved if the initial experts create the items for consensus reaching.(36) In this study, a relevant systematic review assisted in guiding development of questions to ask the experts.

Nasa *et al*.(37) described significant variation in how Delphi studies are conducted and reported, which can lead to uncertainty to the conclusions. Using a 9-point qualitative evaluation for a Delphi quality assessment, this study would score 8 out of 9, missing a full score only as group stability could not be evidenced. Transparency of reporting in this study is consistent with all areas in the Conducting and Reporting Delphi Studies (CREDES) checklist(39) and seen in supplementary material.

Sixteen participated in round 1 of a heterogenous panel and although, no minimum number for Delphi method exists, 15 has been suggested as minimum group size.(37) The threshold for consensus was achieved for all measure related questions, which was set *a priori*. The number of rounds of Delphi is dependent on the method and the intended outcomes. In a another modified-Delphi process, exploring learning and assessment in healthcare education, 2 rounds were used to create a consensus.(38) Delphi response rate ideally should not fall below 70% on subsequent rounds(22): this standard was achieved in this study. Quantitative analysis was not extensive in this study due to methodology to summarise items with thematic analysis. This can be considered as a limitation to the study as Kappa values indicated fair agreement only and subsequent rounds were not conducted to seek convergence. Similarly statistical stability could not be tested for the strength of the item responses.(39) Final consensus criteria were set by a third round as an in-person virtual meeting, whereby descriptive statistics of mean ranks complemented by visual box plots to indicate spread, and a thematic map, were available to support the discussion. Thematic analysis may introduce bias, potentially inherent in individual’s interpretations.(40) The primary investigator had a medical educational background, and his and the co-author’s reflexive practice, and being conscious of this in the analysis, aimed to minimise impact of bias of personal experiences. Any measure developed from this process should be tested in different settings, which will reassure with regard to its validity.

The Delphi is not a psychometric instrument in itself(15) but a practical method for gauging a group-based judgment, more useful for complex concepts and decisions. It can be argued that there is not a preferred Delphi method(41), however, the process of a groupthink interaction generates ideas and aids the conceptual framework that this study addressed.

## Conclusion

This study explained the process of sourcing opinion from a varied cohort of experts to determine how the experience of a learner in healthcare practice using immersive technology might be measured. Synthesising these results in to a multidomain instrument requires a systematic approach to testing in future research. There is variation between educationalists in what is regarded as important in learning from this technology, however, it is important to establish relevant experts’ opinions before designing a user experience outcome measure, and this study highlighted a consensus on the topic.

## REFERENCES

1. Aebersold M, Tschannen D, Bathish A. Innovative Simulation Strategies in Education. Nursing Research and Practice. 2012;2012.

2. Hamstra SJ, Brydges R, Hatala R, Zendejas B, Cook DA. Reconsidering Fidelity in Simulation-Based Training. Academic Medicine. 2014;89(3).

3. Jaskiewicz F, Kowalewski D, Starosta K, Cierniak M, Timler D, Jaskiewicz F, et al. Chest compressions quality during sudden cardiac arrest scenario performed in virtual reality A crossover study in a training environment. MEDICINE. 2020;99(48).

4. Jacobs C, Rigby JM. Developing measures of immersion and motivation for learning technologies in healthcare simulation: a pilot study. Journal of Advances in Medical Education & Professionalism. 2022;10(3):163–71.

5. Forrest K, McKimm J. Healthcare simulation at a glance. Hoboken, NJ: John Wiley & Sons; 2019.

6. Zsigmond I, Buhai A, editors. Augmented Reality in Medical Education, an Empirical Study. 21st International Conference on Computational Science and Its Applications (ICCSA); 2021 Sep 13-16; Cagliari, ITALY2021.

7. Skarbez R, Smith M, Whitton MC. Revisiting Milgram and Kishino’s Reality-Virtuality Continuum. Frontiers in Virtual Reality. 2021;2.

8. Henssen D, van den Heuvel L, De Jong G, Vorstenbosch M, Van Walsum AMV, Van den Hurk MM, et al. Neuroanatomy Learning: Augmented Reality vs. Cross-Sections. Anatomical Sciences Education. 2020;13(3):350–62.

9. Stojanovska M, Tingle G, Tan L, Ulrey L, Simonson-Shick S, Mlakar J, et al. Mixed Reality Anatomy Using Microsoft HoloLens and Cadaveric Dissection: A Comparative Effectiveness Study. Medical Science Educator. 2020;30(1):173–8.

10. Tang YM, Chau KY, Kwok APK, Zhu T, Ma X. A systematic review of immersive technology applications for medical practice and education - Trends, application areas, recipients, teaching contents, evaluation methods, and performance. Educational Research Review. 2022;35:100429.

11. Jacobs C. Immersive technology in healthcare education: a scoping review osf.io/tpjyw 2022[

12. Messick S. Validity of psychological assessment: Validation of inferences from persons’ responses and performances as scientific inquiry into score meaning. American Psychologist. 1995;50(9):741–9.

13. Cook DA, Hatala R. Validation of educational assessments: a primer for simulation and beyond. Advances in Simulation. 2016;1(1):31.

14. Gerup J, Soerensen CB, Dieckmann P. Augmented reality and mixed reality for healthcare education beyond surgery: an integrative review. Int J Med Educ. 2020;11:1–18.

15. Belton I, MacDonald A, Wright G, Hamlin I. Improving the practical application of the Delphi method in group-based judgment: A six-step prescription for a well-founded and defensible process. Technological Forecasting and Social Change. 2019;147:72–82.

16. Eubank BH, Mohtadi NG, Lafave MR, Wiley JP, Bois AJ, Boorman RS, et al. Using the modified Delphi method to establish clinical consensus for the diagnosis and treatment of patients with rotator cuff pathology. BMC Medical Research Methodology. 2016;16(1):56.

17. Novakowski N, Wellar B. Using the Delphi Technique in Normative Planning Research: Methodological Design Considerations. Environment and Planning A: Economy and Space. 2008;40(6):1485–500.

18. Dalkey N, Helmer O. An experimental application of the Delphi method to the use of experts. Management science. 1963;9(3):458–67.

19. Keeney S, Hasson F, McKenna HP. A critical review of the Delphi technique as a research methodology for nursing. International Journal of Nursing Studies. 2001;38(2):195–200.

20. de Villiers MR, de Villiers PJT, Kent AP. The Delphi technique in health sciences education research. Medical Teacher. 2005;27(7):639–43.

21. Schneider P, Evaniew N, Rendon JS, McKay P, Randall RL, Turcotte R, et al. Moving forward through consensus: protocol for a modified Delphi approach to determine the top research priorities in the field of orthopaedic oncology. BMJ Open. 2016;6(5):e011780.

22. Kilroy D, Driscoll P. Determination of required anatomical knowledge for clinical practice in emergency medicine: national curriculum planning using a modified Delphi technique. Emergency Medicine Journal. 2006;23(9):693.

23. Franklin KK, Hart JK. Idea Generation and Exploration: Benefits and Limitations of the Policy Delphi Research Method. Innovative Higher Education. 2007;31(4):237–46.

24. Rowe G, Wright G. Expert Opinions in Forecasting: The Role of the Delphi Technique. In: Armstrong JS, editor. Principles of Forecasting: A Handbook for Researchers and Practitioners. Boston, MA: Springer US; 2001. p. 125–44.

25. Jacobs C, Maidwell-Smith A. Learning from 360-degree film in healthcare simulation: a mixed methods pilot. Journal of Visual Communication in Medicine. 2022:1–11.

26. Braun V, Clarke V, Boulton E, Davey L, McEvoy C. The online survey as a qualitative research tool. International Journal of Social Research Methodology. 2020:1–14.

27. Braun V, Clarke V. What can “thematic analysis” offer health and wellbeing researchers? International journal of qualitative studies on health and well-being. 2014;9:26152-.

28. Stevens B, McGrath P, Yamada J, Gibbins S, Beyene J, Breau L, et al. Identification of pain indicators for infants at risk for neurological impairment: A Delphi consensus study. BMC Pediatrics. 2006;6(1):1.

29. Landis JR, Koch GG. The measurement of observer agreement for categorical data. biometrics. 1977:159–74.

30. Boulkedid R, Abdoul H, Loustau M, Sibony O, Alberti C. Using and Reporting the Delphi Method for Selecting Healthcare Quality Indicators: A Systematic Review. PLOS ONE. 2011;6(6):e20476.

31. Catalin Toma IP. The Delphi Technique: Methodological Considerations and the Need for Reporting Guidelines in Medical Journals. International Journal of Public Health Research. 2016;4:47–59.

32. Woodcock T, Adeleke Y, Goeschel C, Pronovost P, Dixon-Woods M. A modified Delphi study to identify the features of high quality measurement plans for healthcare improvement projects. BMC Medical Research Methodology. 2020;20(1):8.

33. Suh A, Prophet J. The state of immersive technology research: A literature analysis. Computers in Human Behavior. 2018;86:77–90.

34. Mokkink LB, Terwee CB, Patrick DL, Alonso J, Stratford PW, Knol DL, et al. The COSMIN study reached international consensus on taxonomy, terminology, and definitions of measurement properties for health-related patient-reported outcomes. Journal of Clinical Epidemiology. 2010;63(7):737–45.

35. Cronbach LJ, Meehl PE. Construct validity in psychological tests. Psychological Bulletin. 1955;52:281–302.

36. Iqbal S, Pipon-Young L. The Delphi Method. The Psychologist. 2009;22(7):598–601.

37. Clayton MJ. Delphi: a technique to harness expert opinion for critical decision-making tasks in education. Educational Psychology. 1997;17(4):373–86.

38. Tonni I, Oliver R. A Delphi approach to define learning outcomes and assessment. European Journal of Dental Education. 2013;17(1):e173–e80.

39. Nasa P, Jain R, Juneja D. Delphi methodology in healthcare research: How to decide its appropriateness. 2021 (2222-0682 (Print)).

40. Dodgson JE. Reflexivity in Qualitative Research. Journal of Human Lactation. 2019;35(2):220–2.

41. Boje DM, Murnighan JK. Group Confidence Pressures in Iterative Decisions. Management Science. 1982;28(10):1187–96.

